# De novo design of metalloproteases for targeted amyloid-β cleavage

**DOI:** 10.64898/2026.01.06.697903

**Authors:** Yannan Qu, Chentong Wang, Hongli Zhu, Yanjun Wang, Longxing Cao

## Abstract

De novo protein design has not yet achieved the creation of proteases capable of selectively cleaving any desired peptide bond within a native protein with high precision. Here, we report the use of the flow-based generative model Proteus2 to design metalloproteases by generating enzyme–substrate complexes conditioned on a target peptide sequence and a predefined catalytic motif. Our approach employs a two-step encapsulation strategy to create clamp-like metalloproteases that bind the target peptide in a manner that maximizes substrate sequence specificity. The generative process simultaneously optimizes the precise positioning of the target peptide bond in a catalytically competent configuration and accurately scaffolds the transition state catalytic residues—both essential for specific and efficient catalysis. Using this strategy, we designed zinc metalloproteases targeting three distinct cleavage sites within the aggregation-prone regions of amyloid-β (Aβ), a key pathogenic factor in Alzheimer’s disease. Experimental characterization validated five enzymes, each capable of precise cleavage at the intended sites with high specificity and minimal or undetectable activity on non-cognate substrates. On average, these enzymes accelerated peptide bond hydrolysis by more than 10^7^-fold relative to the uncatalyzed reaction, and enabled efficient digestion of the Aβ peptide into smaller segments when enzymes targeting different sites were combined. Cryo-EM structures of three designed enzymes in complex with Aβ peptide, each targeting a distinct cleavage site, revealed close agreement with the design models. Together, these results demonstrate the potential of sequence-guided, generative approaches for developing programmable, sequence-specific proteolysis and lay the groundwork for future applications in basic research and therapeutic development.

## Introduction

Proteases are among biology’s most powerful regulators^1,2^. By cutting specific peptide bonds they activate hormones^3^, terminate signalling cascades^4^, remodel extracellular matrices^5^, and control protein homostasis^1^. Their catalytic activity also underpins a wide spectrum of industrial and medical applications—from laundry detergents and food processing to wound debridement and thrombolytic therapy^6,7^. Yet the utility of natural proteases is circumscribed by their hard-wired substrate preferences^8,9^. An enduring objective is therefore to create “molecular scalpels” whose cleavage site can be programmed at will, enabling precise dissection of cellular pathways and highly selective therapeutic intervention^10,11^. Despite decades of effort, very few platform delivers custom proteases that combine robust catalytic efficiency with single-bond specificity^12–14^.

Metalloproteases are prime candidates for such re-engineering, and their catalytic chemistry has been well studied^15–17^. In a typical zinc metalloprotease, the zinc ion is coordinated by three amino-acid ligands and positioned adjacent to a general base that activates a water molecule for nucleophilic attack on the peptide bond, while a neighbouring residue stabilises the tetrahedral oxyanion intermediate^17^ (**Fig. 1a,b**). This minimalist engine is embedded in diverse scaffolds—including matrix metalloproteinases^18^, ADAMs^19^, astacins^20^, thermolysins^21^, and bacterial virulence factors^22^—that collectively drive processes from extracellular-matrix turnover to cytokine maturation. Despite their natural versatility, repurposing these enzymes for novel targets remains challenging. While efforts to reprogram specificity through directed evolution or rational mutagenesis have yielded promising advances^12,13,23^, certain structural and functional limitations persist. As a result, repurposed natural metalloproteases have achieved only modest improvements in selectivity and therapeutic potential^24^.

**Fig. 1:**
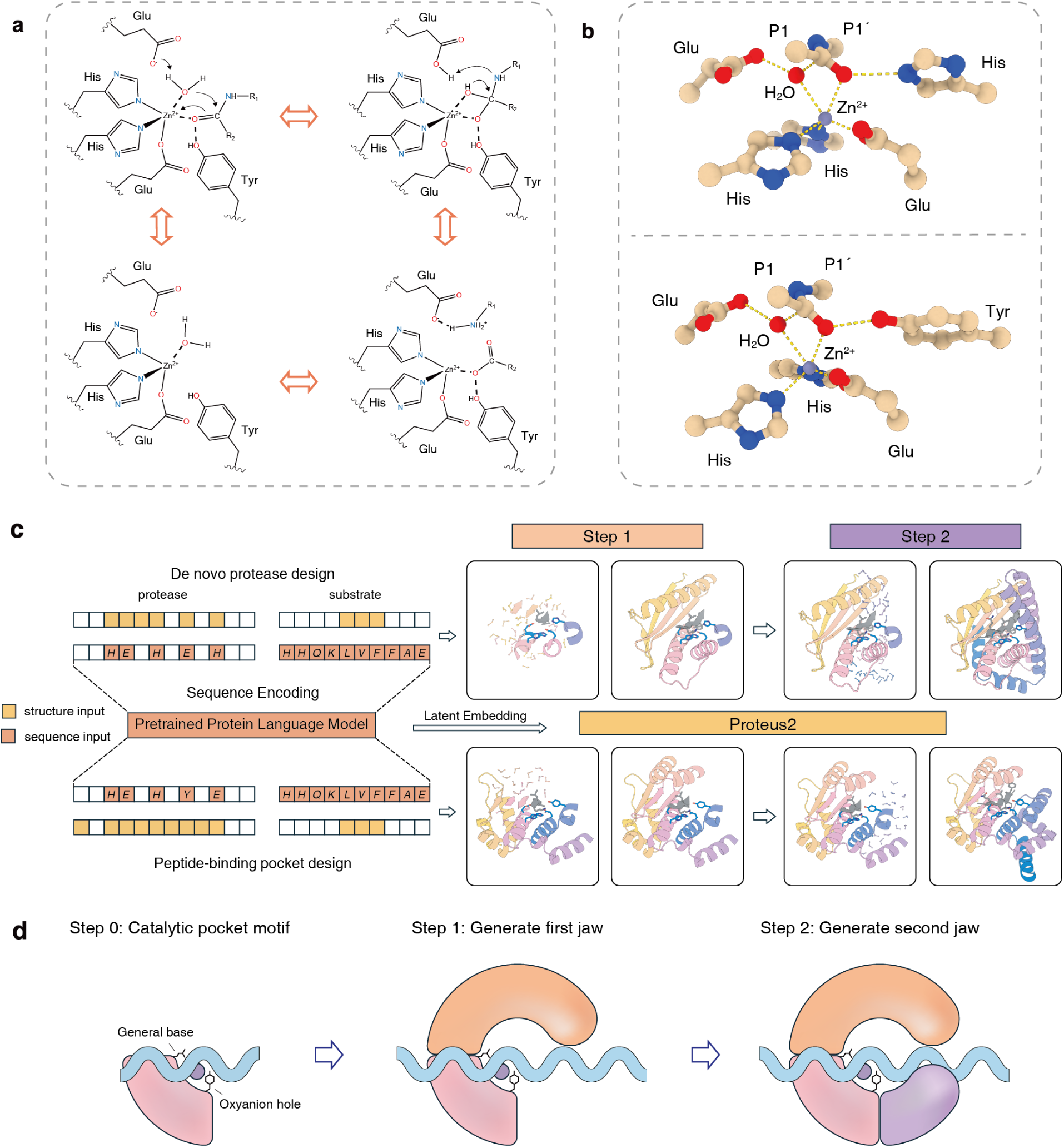
Catalytic mechanism of zinc-dependent metalloproteases and two-step de novo design of clamp-like protease scaffolds. **a,** The catalytic cycle of zinc-dependent metalloproteases involves a precisely orchestrated sequence of molecular events at the active site. The reaction initiates as the scissile carbonyl oxygen of the substrate coordinates with the catalytic zinc ion, polarizing the carbonyl and increasing its electrophilicity. Concurrently, a conserved glutamate residue acts as a general base, deprotonating a zinc-bound water molecule to trigger nucleophilic attack on the carbonyl carbon. This generates a transient tetrahedral intermediate, with its negatively charged oxyanion stabilized via hydrogen bonds within the oxyanion hole—typically formed by a tyrosine or histidine. The catalytic glutamate then functions as a general acid, transferring a proton to the amide nitrogen of the leaving group, thereby facilitating intermediate collapse and peptide bond cleavage. The hydrolyzed products subsequently dissociate, and a new water molecule occupies the zinc coordination sphere, restoring the enzyme for further catalytic cycles. **b,** Theozyme models for the Zn(II)-activated water nucleophilic attack on the substrate peptide bond. Both panels illustrate the core catalytic components: the central Zn(II) ion coordinated by three endogenous amino acid residues, the catalytic glutamate acting as a general base, the nucleophilic water molecule, and the substrate scissile bond represented by the P1 and P1’ residues. **c**, Computational framework for designing substrate-specific metalloproteases. Generation is initialized with either a minimalist catalytic motif (de novo design) or a largely preserved template topology (pocket design). A pretrained protein language model encodes the sequences of the substrate and catalytic residues into latent embeddings. These embeddings, alongside the structural motif, condition the Proteus2 generative model for substrate-sequence-aware protease design via a two-step encapsulation strategy. **d**, Schematic representation of the two-step encapsulation strategy. The scaffold is assembled in a sequential manner to achieve complete substrate encapsulation, ensuring that the clamp precisely engages the target peptide for sequence-specific recognition.

De novo protein design holds the promise of sidestepping the intrinsic limitations of natural enzymes by constructing entirely new scaffolds around a chosen chemical task^25–29^. Recent breakthroughs in deep-learning based structure generation have indeed delivered artificial metallohydrolases whose catalytic efficiencies approach those of native enzymes^26,28,30^; however, these successes have been confined almost exclusively to model reactions such as the hydrolysis of activated ester bonds in small, hydrophobic substrates—reactions that present relatively low activation barriers and low specificity demands. Extending this progress to true proteolysis is vastly more challenging. Amide bonds are orders of magnitude more inert than esters^31,32^, and a protease must not only accelerate their cleavage but also capture a highly flexible, polar peptide and orient one particular bond in a near-attack conformation. Moreover, practical applications require a high level of sequence selectivity, such that the designed substrate binding groove preferentially accommodates the target sequence while minimizing activity toward closely related peptides. Natural proteases achieve this balance with folds that have been evolutionarily honed for cognate substrates; even slight structural perturbations can disrupt the sub-angstrom precision of the catalytic residues^9,33^. Consequently, conventional sequence redesign has limited success in retargets these enzymes while maintaining native-like catalytic efficiency^33–36^. The central challenge, therefore, is to design new structures that simultaneously preserve the strict metal-dependent catalytic geometry, lock the scissile bond into position, and rigorously discriminate against off-target sequences—an objective that has so far remained out of reach.

### Computational strategy for designing sequence specific metalloprotease

We set out to develop a general strategy for the de novo design of sequence-specific metalloproteases. To establish catalytic motifs for design, we focused on two representative zinc-dependent endopeptidases: PPEP-1 (PDB: 6r4w) ^37^ and Fungalysin (PDB: 4k90) ^38^. Both enzymes are secreted by microorganisms and play critical roles in survival or pathogenicity by hydrolyzing extracellular host or cell-surface proteins. Despite sharing a common catalytic mechanism, these two enzymes exhibit distinct active site architectures. Notably, their oxyanion hole compositions differ: PPEP-1 employs a tyrosine, whereas Fungalysin utilizes a histidine residue. Structurally, PPEP-1 features a broad, clamp-like groove formed by two flexible loops that close over the substrate, conferring high sequence specificity for cleavage at Pro-Pro amide bonds. To reprogram this specificity, we developed a strategy that removes the peptide-binding loops from PPEP-1, while retaining the core protein structure (**Fig. 1c**). The excised regions were then rebuilt, enabling the design of new peptide-binding interfaces. Proteases designed through this approach are denoted “PP” (peptide-binding pocket designed protease) followed by a numerical identifier. In contrast, Fungalysin lacks an extended, pre-formed substrate-binding pocket and instead features an open substrate binding groove, resulting in a broader substrate specificity for peptide bonds flanked by hydrophobic side chains. For this scaffold, we extracted only the minimalist catalytic motif—comprising the zinc-coordinating residues, the general base glutamate, and the oxyanion hole histidine (**Fig. 1c**). This minimalist approach necessitates de novo generation of the entire surrounding protein scaffold to accurately position the catalytic motif and design a novel peptide-binding interface for sequence-specific recognition. Proteases generated using this minimalist catalytic motif are designated “DP” (de novo protease).

To enable precise co-design of enzyme functionality and substrate specificity, we employed Proteus2, our generative flow-based model significantly advanced over its predecessor^39^. To explicitly endow the structural generative process with a deep understanding of substrate sequence constraints and specificity, we integrated a customized protein language model pre-trained on UniRef50^40^ with an auxiliary structural decoding objective (see Methods). Proteus2 was trained on experimental structures from the Protein Data Bank^41^ and high-confidence AlphaFold2^42^ predictions using a dual-modality motif scaffolding task. In this framework, sequence and structure conditions are treated as independent masking tracks, leveraging geometry-aware sequence semantics to control structure generation. This decoupled conditioning strategy empowers the model to process hybrid inputs comprising structurally defined catalytic motifs and coordinate-free substrate sequences, thereby generating metalloproteases with stabilized catalytic sites and precise substrate specificity.

We leveraged Proteus2 to co-generate metalloprotease–substrate complex structures using the extracted active motif configurations as templates. To determine the catalytically competent conformation of the substrate relative to the catalytic residues, we used AlphaFold3^43^ to model the substrate peptide configuration within the template protease. The predictions revealed that the substrate adopted a β-strand conformation in both enzymes (**Extended Data Fig. 1**). From these predicted structures, we extracted the P2–P1′ backbone atoms of the Fungalysin substrate peptide (where P1 is the residue immediately preceding, and P1′ immediately following, the cleavage site), and the P3–P1 backbone atoms of the PPEP-1 substrate. In both cases, these segments form strand pairing interactions with the enzyme’s strand adjacent to the active site, thereby enabling the construction of a complete target active site. To maximize structural complementarity and specificity for the substrate peptide, we developed a ‘two-step encapsulation’ strategy based on Proteus2 (**Fig. 1c,d**). Unlike conventional one-step generative approaches—which frequently yield suboptimal packing and limited control over overall enzyme topology—our method proceeds sequentially. The first step generates the protein backbone surrounding the catalytic glutamate, stabilizing the general base and achieving partial pre-wrapping of the substrate. In the second step, the scaffold is constructed around the oxyanion hole to complete substrate peptide encapsulation. This stepwise process minimizes steric clashes, substantially enhances the packing density at the enzyme–substrate interface, and consistently generates clamp-like enzymes that precisely grasp the substrate peptide—thereby improving sequence-specific recognition. Following backbone generation, protein sequences were optimized using LigandMPNN^44^, and the resulting protease–substrate–zinc complex structures were validated with AlphaFold3 to ensure convergence to the intended catalytic geometry.

### De novo proteases facilitate the direct hydrolysis of Aβ peptide

We selected the amyloid-β (Aβ) peptide as the substrate for design, given its central role in Alzheimer’s disease (AD) pathology, where the aggregation of neurotoxic Aβ plaques is a hallmark event^45^. The pathological self-assembly of Aβ is primarily driven by intermolecular interactions within two hydrophobic regions: the central hydrophobic core (residues 17–21) and the C-terminal segment (residues 30–42) (**Fig. 2a**). These domains act as critical nucleation sites for β-sheet stacking, facilitating the formation of stable, neurotoxic fibrils^46,47^. Consequently, these regions represent key therapeutic targets for catalytic interventions aimed at destabilizing amyloid architecture and mitigating AD progression^48^. To demonstrate the generalizability of our computational methodology, we selected three distinct cleavage sites within Aβ: one within the central hydrophobic core and two within the C-terminal hydrophogic region. We employed both the peptide-binding pocket (PP) and de novo protease (DP) strategies to design metalloproteases targeting each site. On average, 26 designs were selected from each strategy–substrate pair for experimental characterization (**Extended Data Table 2**). These designed enzymes were expressed in *Escherichia coli*, purified using Ni-NTA magnetic beads, with the His-tag subsequently removed via HRV3C protease. Final purification was performed by size exclusion chromatography. To assess catalytic activity, we engineered fusion substrates in which the targeted Aβ fragment was inserted between the mannose-binding protein (MBP) and a small ubiquitin-like modifier (SUMO) tag. Successful proteolytic cleavage results in two lower-molecular-weight fragments, readily detectable by SDS-PAGE (**Fig. 2b**). Upon overnight incubation of the designed proteases with their respective substrates in the presence of zinc, we observed that the DP strategy yielded active enzymes for all three targeted sites: DP720-S1, DP622-S2, and DP221-S3. In contrast, the PP strategy generated active designs only for the first site, resulting in PP507-S1 and PP532-S1 (**Fig. 2c**). We attribute the limited success of the PP designs at the remaining sites to steric constraints imposed by excessive preservation of the template backbone, which may be incompatible with the sequence and geometry of these cleavage sites. In contrast, the DP strategy allows broader sampling of the conformational space, facilitating the discovery of active variants. Proteolytic specificity was confirmed by mass spectrometry analysis of the cleavage products, with observed molecular weights matching the theoretical predictions and validating site-specific cleavage (**Extended Data Fig. 2**). Notably, for DP221-S3, in addition to the major mass spectrometric peak corresponding to the intended cleavage products, a very minor peak—representing approximately 10% of the main product’s abundance—was also observed, indicating an alternative cleavage site shifted by one residue toward the C-terminus (**Extended Data Fig. 2e**). We next characterized their catalytic efficiency by monitoring the time course of substrate cleavage (**Extended Data Fig. 3**). At a 1:2 enzyme-to-substrate molar ratio, DP622-S2 exhibited the highest activity, achieving complete substrate degradation within 4 hours (**Extended Data Fig. 3b**). DP720-S1 and PP507-S1 achieved full digestion in 8 hours (**Extended Data Fig. 3a,c**), while PP532-S1 required nearly 12 hours for complete cleavage (**Extended Data Fig. 3e**). In contrast, DP221-S3 displayed slower kinetics, cleaving more than half of the substrate after 12 hours (**Extended Data Fig. 3d**). Collectively, these results indicate our design strategies enables the de novo generation of sequence-specific metalloproteases capable of targeting disease-relevant sites within Aβ.

**Fig. 2:**
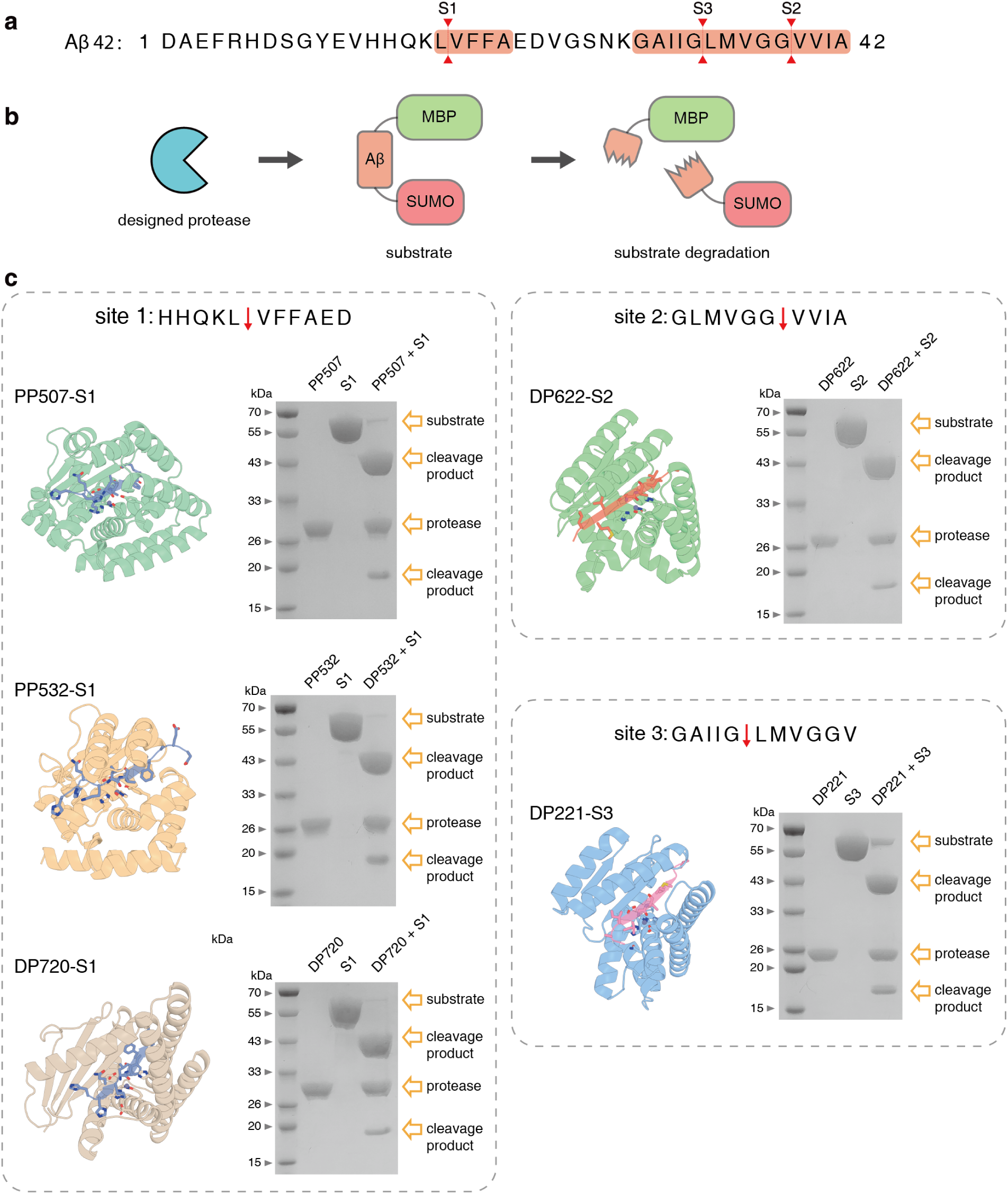
Experimental validation of designed metalloproteases targeting Aβ. **a**, Primary sequence of Aβ42. The complete amino acid sequence of Aβ42 is shown, with two main hydrophobic domains highlighted in orange and three selected cleavage sites marked in red. **b**, Schematic of the proteolysis assay. To facilitate expression and protelytic analysis, Aβ segment was engineered as a fusion protein (MBP-Aβ_segment-SUMO). The diagram illustrates that specific cleavage within the Aβ sequence by the designed proteases results in separation of the MBP and SUMO fragments, which are readily detected via molecular weight shifts in SDS-PAGE experiments. **c**, Design models and proteolytic screening of designed proteases. Design models of five designed proteases are displayed alongside SDS-PAGE analysis, which confirms the successful cleavage of the MBP-Aβ-SUMO substrate.

### Validation of catalytic mechanism

To elucidate the catalytic mechanism and confirm the functional roles of key active site residues, we conducted a series of site-directed mutagenesis experiments (**Fig. 3a**). For designs generated by both the DP and PP strategies, the catalytic glutamate and oxyanion hole residues (histidine in the DP designs and tyrosine in the PP designs) were mutated to alanine (**Fig. 3b**). Additionally, in the DP designs, we mutated the newly generated second-sphere residues that form hydrogen bonds with the zinc-coordinating residues, to evaluate their contribution to catalytic activity. Functional assays revealed that substitution of the catalytic glutamate with alanine universally abolished proteolytic activity across both DP and PP designs, highlighting its indispensable role as the general base in catalysis (**Fig. 3b**). Interestingly, in the DP designs, mutation of the oxyanion hole histidine to alanine had minimal or even positive effects on protease function; notably, the H191A variant of DP221-S3 exhibited enhanced substrate degradation. By contrast, in the PP designs, alanine substitution of the oxyanion hole tyrosine resulted in partial reduction of activity. These findings suggest that the oxyanion stabilization function may be partially compensated by solvent molecules in the absence of the native residue. Furthermore, mutagenesis of the second coordination sphere residues in the DP designs underscored their importance in tuning catalytic efficiency. The D128A mutation in DP720-S1 and the N111A mutation in DP221-S3 each resulted in a marked reduction in activity. In DP622-S2, single mutations (D126A or Y91F) had minimal impact on catalytic activity, whereas the double mutant (D126A/Y91F) led to a dramatic decrease in activity (**Fig. 3b**). These results suggest that second-shell interactions play context-dependent roles in stabilizing the primary coordination environment and modulating the electrostatic landscape of the active site. Collectively, these findings support the conclusion that the observed catalytic activity arises from the designed active sites and highlight the importance of precise active site architecture for efficient catalysis.

**Fig. 3:**
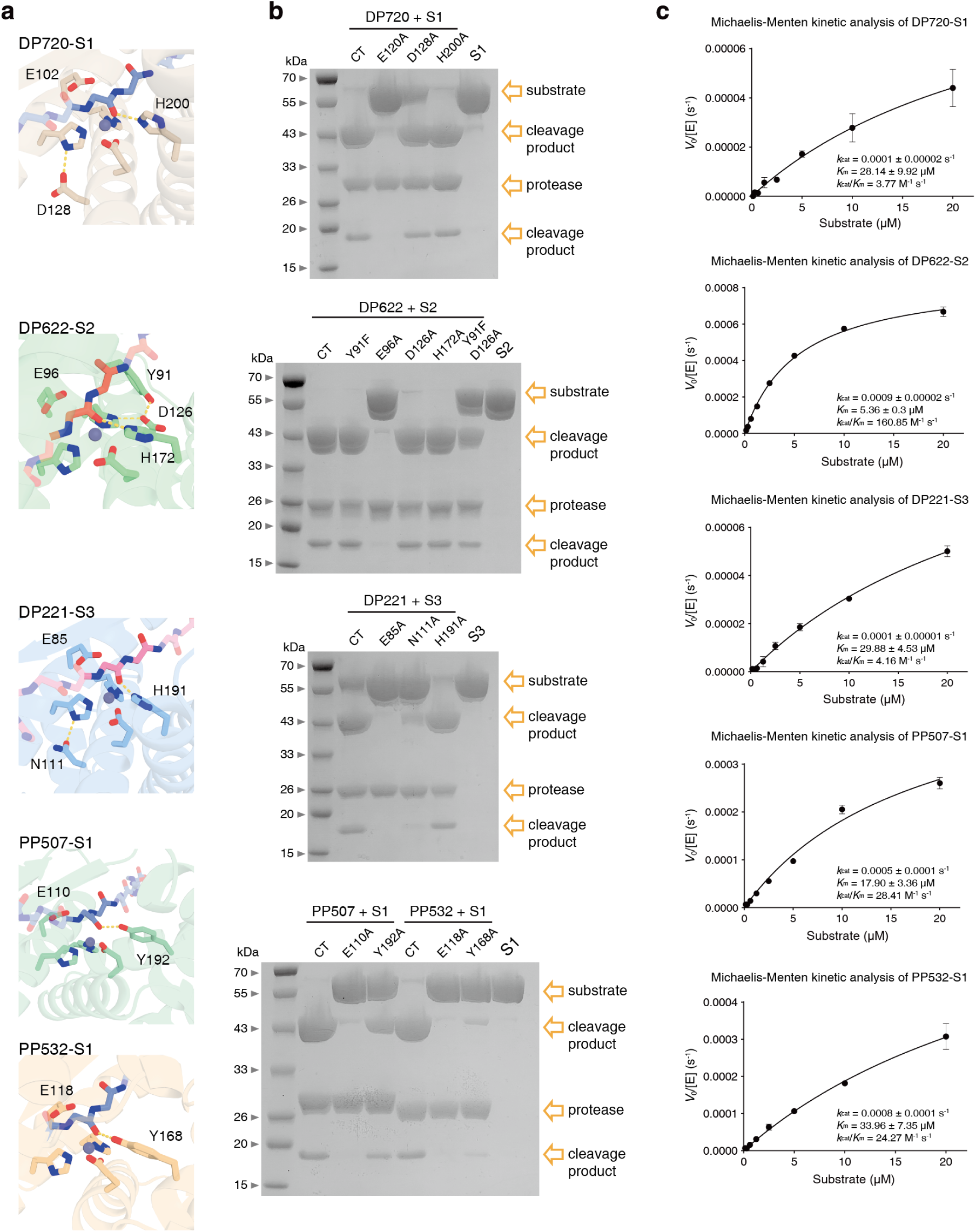
Mutagenesis and kinetic characterization of the designed metalloproteases. **a,** Atomic details of the active site architectures in the design models, highlighting the precise spatial arrangement of key catalytic residues. The coordination of the zinc ion by histidines and glutamates, the positioning of the general base, and the configuration of the oxyanion hole are illustrated. **b,** Mutational analysis of catalytic residues. SDS-PAGE analysis of various catalytic residue mutants is shown, verifying the functional roles of the designed active site residues. **c,** Kinetic characterization of the designed proteases. Proteolytic kinetics were assessed by measuring initial reaction velocities at varying substrate concentrations. Data were fitted to the Michaelis-Menten equation to determine key kinetic parameters, including the Michaelis constant (*K_m_*) and turnover number (*k_cat_*). The calculated *k_cat_*/*K_m_* values quantify the catalytic efficiency of the de novo designed enzymes, demonstrating their substrate recognition and cleavage ability. Data represent mean ± standard deviation (black dots and error bars).

### Characterization of the catalytic kinetics

To quantitatively assess the kinetics of our designed proteases, we developed a specialized fluorescence-based assay. The intrinsic hydrophobicity of the Aβ target sequences posed significant challenges for synthesizing peptides compatible with standard fluorophore-quencher systems, due to marked solubility and aggregation issues. As an alternative, we engineered a FRET-based biosensor by inserting the targeted Aβ sequences directly between the fluorescent protein pair mTurquoise2 (donor) and mVenus (acceptor) (**Extended Data Table 3**). In this system, enzymatic cleavage of the substrate separates the fluorescent proteins, resulting in a measurable decrease in FRET efficiency. Kinetic parameters were determined by fitting initial reaction velocities to the Michaelis-Menten model (**Fig. 3c**). Analysis revealed that the turnover numbers (*k_cat_*) for our designs were generally on the order of 10^-4^ s^-1^, with Michaelis constants (*K_m_*) in the micromolar range. These *k_cat_* values represent an average > 10^7^-fold rate enhancement relative to the uncatalyzed hydrolysis of peptide bonds^31^. Among the designed proteases, DP622-S2 demonstrated the highest catalytic efficiency, achieving a *k_cat_*/*K_m_* of 160.8 M^-1^s^-1^, driven by a favorable combination of a high *k_cat_* (0.0009 s^-1^) and a low *K_m_* (5.36 μM). The PP507-S1 and PP532-S1 exhibited moderate efficiencies of 28.4 and 24.3 M^-1^s^-1^, respectively, while DP221-S3 and DP720-S1 displayed the lowest efficiencies at 4.2 and 3.8 M^-1^s^-1^. With the exception of DP720-S1, these kinetic findings were in close concordance with our time-course degradation assays. The discrepancy observed for DP720-S1—which showed effective substrate degradation comparable to PP507-S1 in previous assays but exhibited low catalytic efficiency in the FRET system—may be attributable to steric hindrance, as the bulky fluorescent proteins could restrict substrate access to the active site. Notably, the *K_m_* value of our lead design, DP622-S2, closely parallels those of naturally occurring Aβ-degrading proteases such as Insulin-Degrading Enzyme, which also exhibit *K_m_* values in the low micromolar range^49^. Although the turnover rates of our designs are lower than those of natural enzymes, achieving micromolar substrate affinity demonstrates that our computational strategy can successfully encode robust recognition for disease-relevant targets.

To further elucidate the functional roles of the oxyanion hole and second coordination sphere residues, we characterized the catalytic kinetics of the corresponding mutants, including single mutants of DP622-S2 (Y91F, D126A, or H172A), DP221-S3 (H191A), as well as the DP622-S2 double mutant (Y91F/D126A) (**Extended Data Fig. 4**). Intriguingly, all tested single mutations in the DP622-S2 series (Y91F, D126A, and H172A) (**Extended Data Fig. 4a-c**) showed enhanced catalytic efficiency (*k_cat_*/*K_m_*) compared to the parental design, whereas the double mutant Y91F/D126A decreased catalytic efficiency (**Extended Data Fig. 4d**), consistent with the SDS-PAGE results. The observed improvements were predominantly driven by increased turnover numbers (*k_cat_*); notably, the DP622-S2 D126A variant exhibited a threefold increase in *k_cat_* (**Extended Data Fig. 4b**). This suggests that optimizing the catalytic microenvironment can significantly modulate the activation energy barrier of the rate-limiting step. Most mutations led to a modest decrease in substrate affinity, as reflected by an increase in *K_m_*; however, the DP622-S2 Y91F variant maintained a *K_m_* nearly identical to the original, resulting in a twofold enhancement in overall catalytic efficiency **(Extended Data Fig. 4a)**. The DP221-S3 H191A mutation had a synergistic effect, increasing *k_cat_* while simultaneously decreasing *K_m_*, resulting in a 2.7-fold improvement in catalytic efficiency **(Extended Data Fig. 4e)**. Together, these findings indicate that second-shell coordination and oxyanion hole residues have a substantial impact on catalytic efficiency, largely by modulating *k_cat_*—a parameter that remains challenging to accurately model in current design protocols.

### Characterization the substrate specificity

We next assessed the substrate specificity of the designed metalloproteases by evaluating their cleavage activity against three distinct MBP–Aβ_segment–SUMO fusion constructs (Substrates 1, 2, and 3) (**Fig. 4**). SDS-PAGE analysis of the cleavaged products revealed that PP507-S1 exhibited the highest degree of selectivity, catalyzing cleavage exclusively of Substrate 1, with no detectable activity against the other two substrates (**Fig. 4b**). In contrast, PP532-S1 displayed broader substrate tolerance, efficiently processing all three substrates (**Fig. 4c**), while PP720-S1 showed measurable, though reduced, activity against Substrate 2 (**Fig. 4a**). DP622-S2 and DP221-S3 exhibited partial cross-reactivity with each other’s substrates, albeit at considerably lower efficiencies compared to their cognate targets (**Fig. 4d,e**). We speculate that the observed cross-reactivity for DP622-S2 and DP221-S3 may be attributed to extensive sequence overlap between the target sequences and the use of glycine-serine (GS) linkers flanking the inserted Aβ segments, which may facilitate recognition of similar sequence motifs across different constructs (**Extended Data Table 1**). Notably, the flanking sequence of the S1 cleavage site contains a high proportion of bulky aromatic and large hydrophilic residues, whereas the regions flanking the S2 and S3 sites are primarily composed of small aliphatic amino acids and glycines. This composition may underlie the high specificity of PP507 and likely prevents the S2/S3 enzymes from cleaving the S1 substrate due to steric occlusion.

**Fig. 4:**
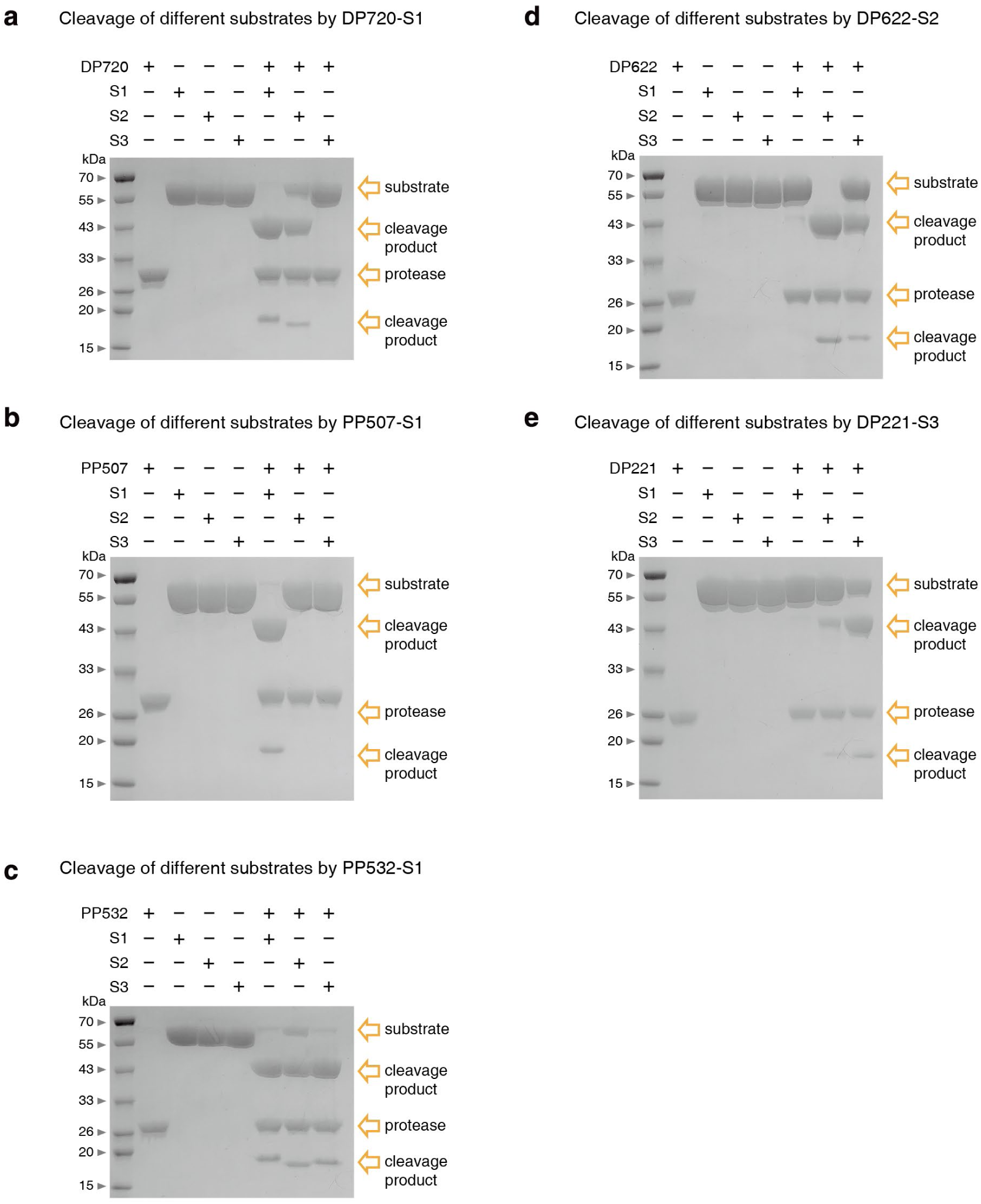
Proteolytic specificity and substrate selectivity of designed proteases. The substrate recognition profiles of five designed proteases (**a**, DP720-S1; **b**, PP507-S1; **c**, PP532-S1; **d**, DP622-S2; **e**, DP221-S3) were systematically evaluated using the three structurally distinct MBP-Aβ_segment-SUMO fusion constructs (Substrates 1, 2, and 3). The results demonstrate the sequence and site selectivity of each protease toward their corresponding substrates.

We next evaluated the substrate specificity of the active site mutants (**Extended Data Fig. 5**). The second-shell and oxyanion hole mutants in DP622 and DP720 exhibited specificity profiles similar to those of their respective parent designs (**Extended Data Fig. 5a,b**). Notably, the DP221-S3 H191A variant showed a remarkable increase in selectivity, virtually abolishing cleavage activity toward non-target substrates (**Extended Data Fig. 5c**). However, mass spectrometry analysis of the cleavage products revealed that the scissile bond shifted almost entirely to the original minor cleavage site—one residue closer to the C-terminus—compared to the parent design (**Extended Data Fig. 6b**). In contrast, the DP622-S2 Y91F mutant retained the intended cleavage site, consistent with the parental enzyme (**Extended Data Fig. 6a**). Nonetheless, the stringent specificity demonstrated by PP507-S1 underscores the potential of our approach to generate custom, sequence-specific proteases. We anticipate that future iterations incorporating explicit negative design strategies during the design and filtering stages will further enhance substrate specificity.

To further evaluate the cleavage activity of our most effective designs and select variants—including PP507, DP622 Y91F, and DP221 H191A, we incubated all three enzymes together with a full-length synthetic Aβ42 peptide. Mass spectrometry analysis revealed a diverse mixture of peptide species. Notably, alongside with several partially cleaved intermediate fragments, we successfully detected the expected fully cleaved products (DAEFRHDSGYEVHHQKL, VFFAEDVGSNKGAIIGL, and VVIA) (**Extended Data Fig. 7**). These results confirm that our de novo metalloproteases are capable of catalyzing the site-specific hydrolysis of the full-length Aβ42 peptide as designed.

### Structural characterization of the protease-substrate complex

To assess the structural accuracy of our computational models, we determined the cryo-EM structures of PP507-S1, DP622-S2, and DP221-S3 in complex with the Aβ42 peptide (**Fig. 5a**). To overcome the resolution limitations inherent to the small molecular weight of the designed proteases, each protease was rigidly fused with a large helical bundle protein generated by Proteus2 (**Extended Data Fig. 8**). Minimal interfacial interactions were computationally introduced to rigidify the fusion construct, ensuring that neither the overall fold nor the active site architecture of the protease was perturbed. To capture the pre-catalytic Aβ42-bound state, the catalytic glutamate in each protease was mutated to glutamine (E to Q). The resulting fusion proteins, with inactivated active sites, were incubated with the Aβ42 peptide and subjected to cryo-EM structure determination (**Extended Data Fig. 9**). Focused refinement of the protease region within the fusion protein yielded structures at resolutions ranging from 2.98 to 3.44 Å (**Extended Data Fig. 10**). The reconstructions revealed clear, continuous density for the Aβ42 substrate, allowing confident assignment of the peptide backbone and local side chains within the binding cleft (**Fig. 5b**). The resolved peptide sequence confirms that all three proteases specifically recognize the designed Aβ segment, consistent with their high substrate specificity.

**Fig. 5:**
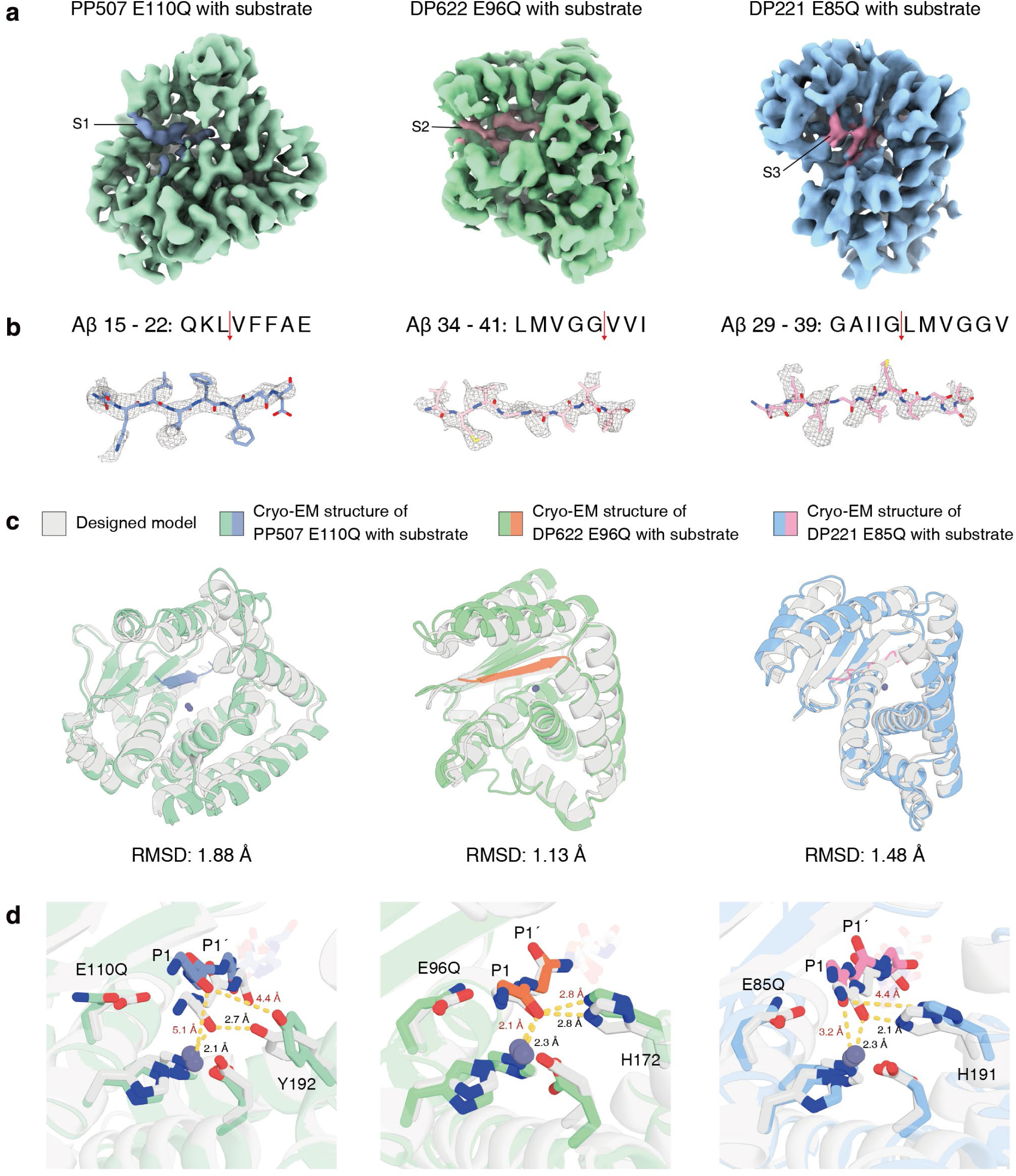
Cryo-EM structure characterization of designed protease in complex with Aβ42. **a,** Overall cryo-EM density maps for each of the three designed proteases. To capture the pre-catalytic substrate-bound complex, the catalytic glutamate in each design was mutated to glutamine (E to Q). **b,** Local cryo-EM density maps highlighting the bound Aβ42 substrate. **c,** Superimposition of the computational design models (grey) and the experimentally determined cryo-EM structures (colored), with backbone Cα RMSD values indicated. **d,** Close-up views of the superimposed active sites. Key catalytic geometries are assessed by measuring the distances between the substrate P1 backbone carbonyl oxygen, the designed oxyanion hole residue, and the catalytic zinc ion. Distance labels are shown in black for the design models and in red for the cryo-EM structures.

Superimposing the experimentally determined cryo-EM structures onto their respective computational models revealed a high degree of global structural agreement, with Cα root-mean-square deviation (RMSD) values ranging from 1.13 to 1.88 Å. This high structural fidelity underscores the robustness of our flow-based generative strategy in accurately constructing the overall protein structures (**Fig. 5c**). Furthermore, detailed inspection of the superimposed active sites confirmed that the designed catalytic geometries were accurately recapitulated in the experimental structures (**Fig. 5d**). Critical spatial arrangements were evaluated by measuring the distances between the substrate P1 backbone carbonyl oxygen, the coordinating zinc ion, and the designed oxyanion hole residue. In PP507-S1 and DP221-S3, we observed local structural relaxation within the binding pocket. For PP507-S1, the P1 oxygen–zinc distance expanded to 5.1 Å in the cryo-EM structure compared to the designed 2.1 Å, and its distance to the tyrosine oxyanion hole increased from 2.7 Å to 4.4 Å. Similarly, DP221-S3 exhibited an enlarged P1 oxygen–zinc distance of 3.2 Å (designed 2.3 Å) and a relaxed distance of 4.4 Å to the histidine oxyanion hole (designed 2.1 Å). In contrast, the active site of DP622-S2 demonstrated remarkable structural fidelity, maintaining a tightly coordinated P1 oxygen–zinc distance of 2.3 Å (closely matching the designed 2.1 Å) and faithfully preserving the designed 2.8 Å distance to the histidine oxyanion hole. The precise geometric pre-organization of the DP622-S2 active site is consistent with our kinetic observations (**Fig. 3c**), which demonstrate both higher substrate affinity and a moderately improved turnover rate for DP622-S2 compared to PP507-S1 and DP221-S3. Collectively, these structural insights, together with biochemical data, underscore that achieving high precision in active site geometry remains critical for enhancing the efficiency of de novo designed enzymes.

## Discussion

By explicitly integrating target sequence information into the flow-based structure generation process, Proteus2 enables the direct generation of sequence-specific metalloproteases. Our two-step encapsulation strategy facilitates the creation of clamp-like proteases that envelop the substrate peptide, thereby maximizing both specificity and discrimination. The successful design of metalloproteases capable of cleaving three distinct native Aβ peptide sequences extends de novo protein design beyond stable binding, enabling functional proteolysis of physiological targets. The atomic-level agreement between the design models and experimentally determined protease-substrate complex structures further demonstrates that our computational approach can achieve both high structure accuracy and precise substrate discrimination. Our specificity characterization also indicates that, for future enzyme design, careful selection of the target site is critical; regions enriched in distinctive side chains, such as bulky aromatic or highly charged residues, may facilitate the development of enzymes with higher specificity. While a certain degree of nonspecific cleavage was seen in some designes, the clamp-like architecture provides a robust starting point for further sequence redesign or structure-guided refinement to enhance specificity.

Our successful de novo design of metalloproteases, starting from minimalist catalytic motifs, represents a major advance in the de novo enzyme design field by bridging the gap between model reactions and catalysis relevant for therapeutic intervention. On average, our designs accelerate peptide bond cleavage by more than 10^7^-fold compared to the uncatalyzed reaction in aqueous solution^31^. Importantly, these metalloproteases exhibit substrate affinities (*K_m_*) comparable to those of natural enzymes, indicating that our design strategies can accommodate the structural and chemical complexity of physiological polypeptides^49^. However, our mutational analysis also indicates that key microenvironmental factors affecting turnover rates (*k_cat_*) remain challenging to accurately model with current approach. While our structural characterization indicates that improving the modeling precision of the active site might represent a promising direction for enhancing catalytic efficiency, further advancements will necessitate careful consideration of solvent effects, second-shell coordination, and the dynamic nature of the catalytic process to achieve optimal catalytic proficiency.

The aggregation of Aβ into neurotoxic plaques is a defining event in Alzheimer’s disease (AD) pathology, making the development of molecular tools to counteract Aβ accumulation a high priority. Current strategies for Aβ clearance, predominantly based on monoclonal antibodies^50^, act by sequestering Aβ for immune-mediated removal rather than direct peptide degradation^51^. In contrast, catalytic degradation offers a compelling alternative, as a single enzyme can process multiple Aβ molecules through direct hydrolysis. Although endogenous proteases like neprilysin and insulin-degrading enzyme can target Aβ^52,53^, their broad specificity presents a risk for undesirable off-target effects when therapeutically enhanced^54,55^. Our work provides a fundamentally distinct strategy—fully de novo design of programmable, sequence-specific proteases for Aβ peptide degradation—enabling the selective hydrolysis of pathogenic sequences and facilitating efficient monomer clearance. By targeting cleavage within the hydrophobic, aggregation-prone regions of Aβ, these proteases have the potential to inhibit the formation of toxic fibrils. Notably, the cleavage site for our most specific design, PP507-S1, is only one residue away from the α-secretase site responsible for the non-pathogenic processing of APP, showcasing the potential for precision engineering^56^. As de novo designed enzymes can target different sites, our multi-enzyme cocktail approach may reveal new possibilities for developing therapeutic interventions for AD.

Overall, our work establishes a robust and generalizable framework for the de novo design of programmable, sequence-specific metalloproteases with wide-ranging applications in research and medicine. Our design approach can be readily adapted to other protease classes, such as serine or cysteine proteases, opening new possibilities for selective protein manipulation and targeted protein degradation that are unattainable with naturally occurring enzymes. Looking ahead, the integration of explicit negative design strategies, machine learning-guided optimization, and advanced transition-state modeling will likely yield next-generation proteases with even higher specificity and catalytic efficiency. Ultimately, this technology sets the stage for the rational development of “designer” proteases for a broad spectrum of diseases and empowers both basic research and therapeutic innovation by providing unprecedented control over protein function in living systems.

## Data availability

The density maps and structure files have been deposited into the Electron Microscopy Data Bank and the RCSB Protein Data Bank with the following accession codes: EMD-69321 and 23WM (PP507 E110Q); EMD-69322 and 23WN (DP622 E96Q); EMD-69323 and 23WO (DP221 E85Q).

## Code availability

The Rosetta macromolecular modelling suite (https://www.rosettacommons.org) is freely available to academic and commercial users. Design protocols and analysis scripts used in this paper are available in the Methods and at xxx. The code for Proteus2 is currently uploaded to Zenodo at xxx.

## Acknowledgements

We extend our sincere thanks to the Dang Lab at Westlake University for their valuable guidance and technical expertise in peptide synthesis. We would like to thank the Mass Spectrometry Facility of Westlake University for help in sample analysis; and the Westlake University HPC Center for computation assistance.

## Author contributions

Y.Q., C.W. and L.C. designed the research. Y.Q., C.W. and L.C. developed the design pipeline and made the designs. Y.Q., H.Z. and Y.W. pefromed the experiments. All authors analysed data. L.C. supervised research. Y.Q., C.W. and L.C. wrote the manuscript with the input from the other authors. All authors revised the manuscript.

## Competiting interests

A provisional patent application will be filed prior to publication.

## Funding sources

This work was supported by grants from the National Key R&D Program of China (2022YFA1303700), the National Natural Science Foundation of China (32370989, 3257120243), The Zhejiang Province Pioneer Plan (2025C01189), and Westlake Education Foundation to L.C.

## Notes

### Competing Interest Statement

The authors have declared no competing interest.

### Summary of Updates

Some formatting errors were corrected, and several redundant figures were removed.

